# Ethanol withdrawal induces temporally distinct alterations in oxytocinergic, opioidergic, glutamatergic and neuroimmune systems associated with recognition memory deficits

**DOI:** 10.64898/2026.09.19.752895

**Authors:** Sydney A Zimmerman, Anna Onisiforou, Panos Zanos, Maria Sklirou, Juan-Antonio Garcia Carmona, Fani Pantouli, Lisa Wells, Alexis Bailey, Polymnia Georgiou

## Abstract

Alcohol use disorder is associated with persistent neuroadaptations and cognitive impairments, particularly during withdrawal, yet the temporal dynamics and interactions between these processes remain poorly understood. In the present study, we investigated time-dependent alterations in the oxytocin receptor (OTR), μ-opioid receptor (MOPr), metabotropic glutamate receptor 5 (mGlu5R), and translocator protein (TSPO) following chronic ethanol exposure and withdrawal, and their relationship to behavioral impairments. Male C57BL/6J mice were exposed to a 10-day escalating-dose ethanol liquid diet paradigm followed by 1, 4, or 7 days of withdrawal. Motor coordination and recognition memory were assessed using rotarod and novel object recognition tests, and receptor binding was quantified using autoradiography across multiple brain regions. Chronic ethanol exposure induced motor deficits that persisted during early withdrawal, whereas recognition memory impairments emerged after 7 days of withdrawal. Ethanol exposure and withdrawal produced region- and time-dependent alterations across all systems examined, with increased OTR binding in septal regions and transient reductions in basomedial amygdala OTR binding during intermediate withdrawal that normalized following prolonged withdrawal, transient increases in MOPr binding in striatal regions at 4 days of withdrawal, increased mGlu5R binding in the nucleus accumbens core, basolateral amygdala, and hippocampus, and early transient increases in TSPO binding indicative of neuroimmune activation. Notably, mGlu5R binding in the caudate-putamen and amygdala was associated with recognition memory deficits in a withdrawal stage-dependent manner. Bioinformatic analysis of human postmortem hippocampal transcriptomic datasets from individuals with alcohol use disorder revealed no consistent changes in GRM5 expression, suggesting that mGlu5R dysregulation may be driven by post-transcriptional mechanisms. These findings demonstrate that ethanol withdrawal is characterized by temporally distinct neuroadaptations across multiple systems and identify mGlu5R as a key correlate of withdrawal-associated memory impairment, highlighting its potential as a translational therapeutic target.

## Introduction

Alcohol use disorder (AUD) is a chronic relapsing disorder characterized by compulsive alcohol consumption, loss of control over drinking, and the emergence of a withdrawal syndrome upon cessation of alcohol intake ^1–3^. Withdrawal is a critical stage of the addiction cycle, during which individuals experience negative affective states, cognitive dysfunction, and heightened relapse vulnerability ^2,4,5^, and these symptoms frequently persist well beyond the acute withdrawal period ^4,6^. Importantly, alcohol withdrawal comprises distinct clinical stages, with acute withdrawal characterized primarily by somatic symptoms, whereas protracted abstinence is associated with persistent cognitive deficits, craving, sleep disturbances, and increased relapse vulnerability ^7,8^. This suggests that chronic alcohol exposure produces long-lasting adaptations in neural circuits involved in reward processing, cognition, stress responsiveness, and behavioral control. While addiction-related neuroadaptations induced by chronic alcohol exposure have been extensively characterized, far less is known about their temporal regulation during withdrawal and recovery. Importantly, the neurochemical mechanisms underlying alcohol withdrawal-related cognitive decline, and how these mechanisms evolve across distinct stages of abstinence, remain poorly understood. Given the dynamic nature of alcohol withdrawal, systematically characterizing the temporal evolution of behavioral and neurochemical adaptations across distinct withdrawal phases is essential for understanding the mechanisms underlying withdrawal-related cognitive impairment and for identifying stage-specific therapeutic targets to improve treatment outcomes. Such knowledge may also facilitate the development of biomarker profiles capable of identifying individuals at risk of persistent cognitive dysfunction or relapse and inform the optimal timing of therapeutic interventions during abstinence.

Multiple neurotransmitter and neuromodulatory systems have independently been implicated in alcohol dependence and withdrawal. In addition to their established roles in alcohol-related behaviours, many of these systems contribute to cognitive processes including learning, memory, and executive function, suggesting that their dysregulation during withdrawal may contribute to the cognitive impairments observed during abstinence. Among these, the oxytocinergic, opioidergic, and glutamatergic systems have received considerable attention because of their roles in alcohol-related behaviours and cognitive function. Pharmacological enhancement of oxytocin receptor signaling reduces alcohol consumption, attenuates withdrawal symptoms, and decreases relapse-like behavior in preclinical models ^9–11^, while clinical studies suggest beneficial effects on craving and withdrawal severity ^9,12,13^. Similarly, the μ-opioid receptor (MOPr) plays a central role in alcohol reward and reinforcement^14–18^, as also evidenced by the efficacy of opioid receptor antagonists such as naltrexone in the treatment of AUD ^19,20^. The metabotropic glutamate receptor subtype 5 (mGlu5R) has also emerged as an important regulator of alcohol-seeking behavior, relapse, and withdrawal-related responses ^21–23^. Crucially, however, these systems have largely been studied in isolation, with most investigations focusing on a single neurobiological system and a single stage of withdrawal. Consequently, little is known about whether these systems undergo coordinated or dissociable adaptations across the course of withdrawal and how such changes may contribute to withdrawal-related cognitive impairment.

Evidence also points to functional interactions between oxytocinergic and opioidergic signaling ^24,25^, while growing data indicate reciprocal modulation between MOPr and mGlu5R pathways ^26–28^. Together, these findings suggest that alterations within individual neurotransmitter systems may not occur in isolation but rather as part of broader neurobiological adaptations during withdrawal. However, because most studies have focused on individual neurotransmitter systems and single withdrawal time points, little is known about the relative timing and regional specificity of alterations in OTR, MOPr, and mGlu5R expression across abstinence. Consequently, it remains unclear whether specific neurochemical adaptations represent potential therapeutic targets during distinct phases of withdrawal or whether they could serve as biomarkers of persistent cognitive dysfunction and relapse vulnerability.

Compounding this complexity, neuroinflammatory processes have emerged as important contributors to alcohol-induced neuropathology. Chronic alcohol exposure activates microglia and promotes neuroimmune signaling throughout the brain^29–31^, contributing to cognitive dysfunction, neuronal injury, and behavioral abnormalities associated with dependence ^29,32^. Neuroimmune activation is particularly relevant to cognitive impairment, as inflammatory processes mediated by glial cells, including microglia and astrocytes, within the hippocampus and other cognition-related brain regions have been implicated in alcohol-induced deficits in learning and memory^31–34^. Translocator protein (TSPO), a widely used marker of glial activation and neuroinflammatory processes, has been proposed as a valuable biomarker of alcohol-induced neuroimmune alterations^32^. Notably, alterations in TSPO expression have been reported in individuals with alcohol dependence, highlighting the potential clinical relevance of neuroimmune mechanisms in AUD^32^. Critically, the oxytocinergic ^35–37^, opioidergic ^38,39^, and glutamatergic ^40–42^ systems have each been independently linked to the regulation of glial activity and neuroinflammatory signaling. This suggests that withdrawal-induced receptor dysregulation and neuroimmune activation may occur in parallel and could potentially influence one another. However, the temporal relationship between neuroimmune activation and neurotransmitter-specific neuroadaptations during withdrawal remains unclear. Clarifying this relationship may help identify optimal therapeutic windows and evaluate the potential utility of neuroimmune markers as predictors of withdrawal severity, cognitive outcome, or relapse vulnerability.

Despite substantial evidence implicating these systems in alcohol-related behaviors, a comprehensive understanding of their temporal regulation during withdrawal and their association with alcohol withdrawal-related cognitive deficits is lacking. In particular, it remains unclear whether changes in oxytocin receptor (OTR), MOPr, mGlu5R, and translocator protein (TSPO; marker of glial activation and neuroinflammatory processes) binding occur simultaneously during withdrawal, whether they exhibit distinct temporal profiles, and whether such alterations are associated with cognitive and behavioral impairments.

Therefore, the present study investigated the effects of chronic ethanol exposure and withdrawal on OTR, MOPr, mGlu5R, and TSPO binding in the mouse brain using quantitative receptor autoradiography. o enhance the translational relevance of these findings, we additionally examined GRM5 gene expression in postmortem brain tissue from individuals with alcohol use disorder and matched controls. To capture the temporal evolution of these neuroadaptations, receptor binding was examined following chronic ethanol exposure and after 1, 4, and 7 days of withdrawal, representing acute, intermediate, and protracted stages of abstinence, respectively. Behavioral assessments of locomotor activity, motor coordination, and recognition memory were conducted in parallel to determine whether neurochemical changes were associated with withdrawal-related cognitive and functional deficits. Finally, correlation analyses were performed to determine whether alterations in OTR, MOPr, mGlu5R, and TSPO binding were associated with one another and with measures of recognition memory, and motor coordination, thereby identifying neurochemical adaptations associated with withdrawal-related cognitive impairment and functional deficits across the withdrawal period.

## Materials and Methods

### Animal welfare and ethical statement

All animal care and experimental procedures were conducted in accordance with the U.K. Animal Scientific Procedures Act (1986). Animal studies are reported in compliance with the ARRIVE guidelines.

Male C57BL/6J mice (8-week old at the beginning of the experiments, Charles River, UK), were housed individually (Optimice cages with direct exhaust ventilation technology; outside cage dimensions: 34.3 cm L x 29.2 cm W (front) x 15.5 cm H; Animal Care Systems Inc., Colorado, USA) in a temperature-controlled environment with a 12-hour light/dark cycle (lights on: 06:00 am). Only male mice were used in the current study, as epidemiological evidence indicates that men exhibit higher rates of alcohol consumption and alcohol use disorder compared with women ^43^. This sex difference has been consistently reported in large population-based studies, although it is important to note that sex-specific mechanisms in alcohol dependence remain an important area for future investigation. Food and water were available *ad libitum*. Four days prior to the start of the experiment all mice were provided with a combination of liquid diet (Ensure plus, Abbot laboratories, UK) and chow *ad libitum* for two days, followed by exposure to solely liquid diet *ad libitum* for two days to acclimate them to this form of feeding. A mouse model was used in this study as it is commonly used to assess the neurobiological mechanisms underpinning ethanol addiction.

## Materials

Maltodextrin was purchased from Sigma-Alldrich (Poole, UK). Ethanol (96%) was purchased from Fisher Scientific (Loughborough, UK). [^3^H]-DAMGO (specific activity 1905.5 GBq·mmol^−1^) and [^125^I]-OVTA (specific activity 81.4 TBq·mmol^−1^) were purchased from PerkinElmer (Waltham, MA, USA). [^3^H]MPEP (specific activity 2.22 TBq/mmol) was purchased from American Radiolabeled Chemical (St. Louis, MO, USA). [3H]PBR28 (specific activity: 3.03 TBq·mmol^-1^) was custom labelled by Tritech, Switzerland. Unlabelled PK11195 and naloxone was purchased from Sigma-Aldrich (Poole, UK). Fenobam and unlabeled (Thr4, Gly7)-oxytocin were purchased from Tocris Bioscience (Bristol, UK) and Bachem (Bubendorf, Switzerland) respectively.

### Chronic escalating-dose administration paradigm

The Lieber-DeCarli liquid diet containing ethanol paradigm was as previously described with minor modifications^44^. Mice were either exposed to a 10-day escalating dose ethanol-containing diet (2.3% - 2 days, 4.7% - 2 days and 7% - 6 days) or control diet, where the ethanol was isocalorically substituted with maltodextrin (Sigma-Alldrich, Poole, UK). Ethanol-containing and control diets were prepared fresh twice per day (9:00am and 5:30pm). In different cohort of animals (withdrawal groups), following 10 days of ethanol consumption, ethanol was isocalorically replaced with maltodextrin and the mice were withdrawn for one, four or seven days. Matched controls were included (8 groups in total). Water was available *ad libitum* throughout the ethanol administration paradigm and withdrawal. The liquid diet was the only source of food provided to the mice during the ethanol administration paradigm and withdrawal. Body weight, water, food and alcohol consumption was measured daily throughout the length of the experiment. Animals were euthanized by a 30-sec exposure to CO_2_ followed by decapitation 10 days following ethanol exposure (chronic group) and one, four or seven days after the ethanol removal (withdrawal group). Brains were stored at −80°C until use for quantitative receptor autoradiography.

### Behavioral paradigms

All mice underwent behavioral testing. Mice were assessed for locomotor changes, memory impairments and motor coordination problems following the chronic ethanol consumption and following one, four and seven days of withdrawal. Experimental timeline is provided in **Figure 1**.

**Figure 1.**
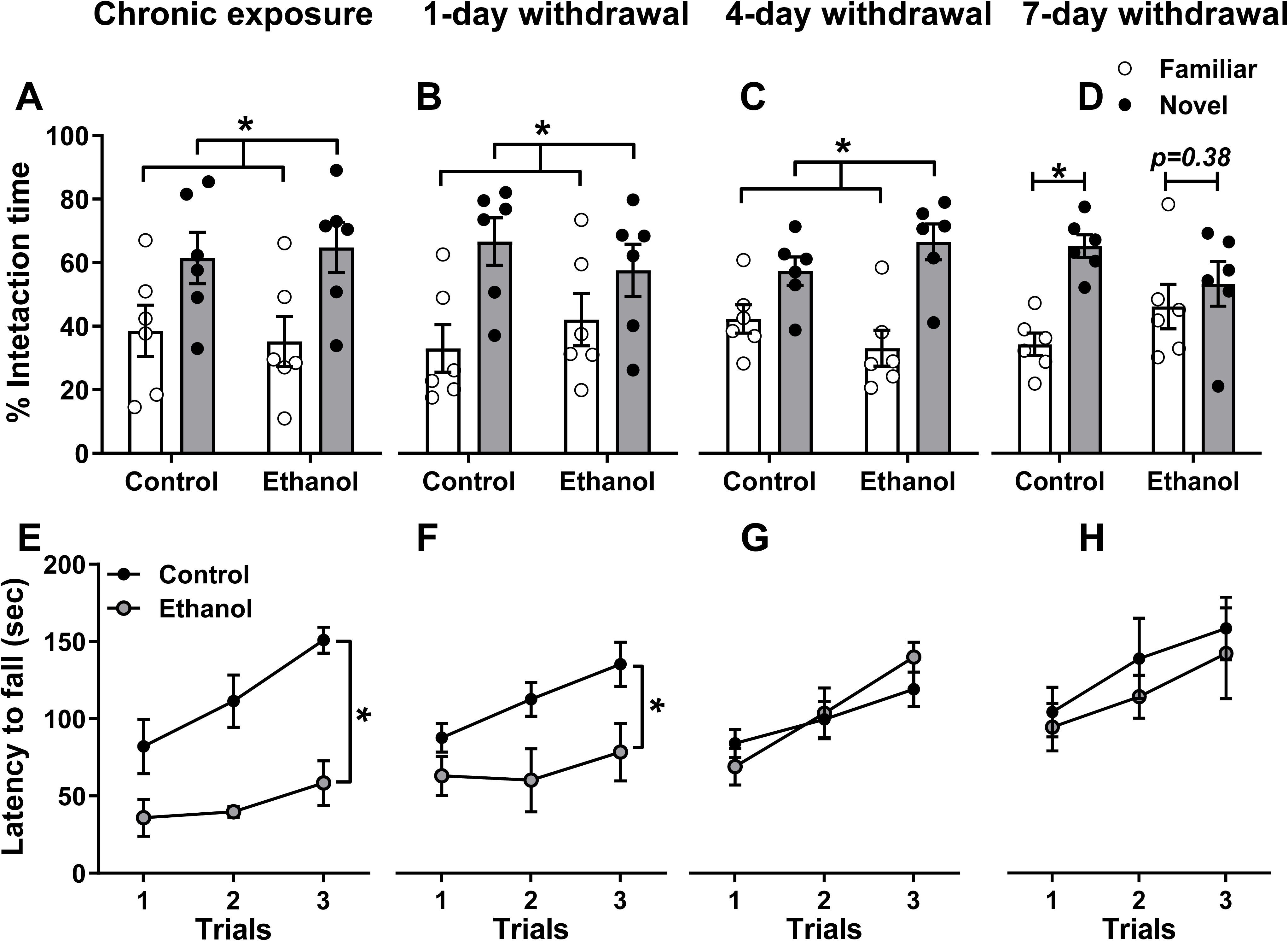
Effects of chronic ethanol exposure and withdrawal on recognition memory and motor coordination. Male C57BL/6J mice were exposed to an ethanol-containing liquid diet for 10 days and subsequently underwent 1, 4, or 7 days of withdrawal. **(A-D)** Novel object recognition performance following chronic ethanol exposure and after 1, 4, and 7 days of withdrawal relative to their respective control groups. **(E-H)** Rotarod performance, expressed as latency to fall, following chronic ethanol exposure and after 1, 4, and 7 days of withdrawal relative to their respective control groups. Data are presented as mean ± SEM (n = 5-6 animals per group). \**p* < 0.05.

Tests were performed from the least stressful to the most stressful. Two hours following the completion of the behavioral experiments Animals were euthanized by a 30-sec exposure to CO_2_ followed by decapitation 1 hour 10 days after the introduction of ethanol (chronic group) and one, four or seven days after the ethanol removal (withdrawal group). Brains were stored at −80°C until use Quantitative Receptor Autoradiography.

#### Locomotor activity

Locomotor activity for each mouse was measured in locomotor chambers (40cm x 20cm x 20cm; Linton Instrumentation, U.K.) for 10 minutes every 2 hours for a total period of 24 hours. For the chronic administration group the first locomotor measurement was taken on day 9, for the 1-day withdrawal group on day 10, for the 4-day withdrawal on day 13 and for the 7-day withdrawal on day 16 at 9am. Distanced travelled and rearing frequency was measured by an automated tracking system (EthoVision v.3.0, Noldus Information Technology, Netherlands).

#### Withdrawal symptoms scoring

During locomotor activity testing, mice were recorded using a digital video-camera (Sony Handycam CX-250, Sony, Japan). Withdrawal symptoms were scored at 0, 2, 4, 8, 12, 16, 20, 24 and 96 hours post-ethanol removal by an experimenter blind to the treatment groups. We did not analyze further than 96 hours as physical withdrawal symptoms have been dissipated but the 96-hour mark. Withdrawal symptoms included backward movements, tremors, grooming and freezing.

#### Novel object recognition

The NOR was performed as previously described ^45,46^ with minor modifications. NOR was performed after locomotion testing. Briefly, the NOR arena used was the same as the one used for the locomotor measures mice to reduce the novelty-induced stress. During the acquisition phase two identical objects (either dice or marbles stuck on a plastic square block) were placed in the arena and mice left to explore both objects for 30 minutes. The order of the objects was alternated between mice to avoid bias towards any of the objects. Following the acquisition phase mice were returned to their home cages for a retention time of 60 minutes. During the testing phase a familiar object and a novel object were placed in the arena and the mouse was re-introduced and left to explore for 2 minutes. All the phases of the NOR were performed on the same day. The sessions were recorded using a digital video-camera (Sony Handycam CX-250, Sony, Japan) and the time spent interacting (directly sniffing) with each object was analyzed using TopScan video tracking software (CleverSys).

#### Rotarod

Rotarod was performed after NOR as it is considered more stressful. Two mice were tested simultaneously on the rotarod (Mouse Rota-Rod 47600, Ugo Basile, Varese, Italy). Once all mice were stationed on the rod, the motor was turned on and the rod was continuously accelerated from 0 rpm at a constant rate of 20 rpm/min over a period of 5 minutes. Mice were returned to their starting position on the static dowel for each consecutive trial (3 trials in total) following a two-minute intra-trial interval. Latency to fall was automatically recorded by the magnetic switch activation.

### Quantitative receptor autoradiography

Coronal brain sections were cut (20 μm thick; 300 μm apart) using a cryostat (Zeiss Microm 505E, U.K.), thaw-mounted onto gelatin subbed ice-cold microscope slides and processed for autoradiography.

#### Oxytocin receptor binding

OTR autoradiography was carried out as previously described^47^. For the determination of total binding, slides were incubated in 50pM [^125^I]-ornithine vasotocin (OVTA) (PerkinElmer, 81.4 TBq/mmol) in an incubation buffer medium (50mM Tris–HCl, 10 mM MgCl_2_, 1mM EDTA, 0.1% w/v bovine serum albumin, 0.05% w/v bacitracin; pH 7.4 at room temperature) for 60 minutes. For the determination of NSB, adjacent sections were incubated with [^125^I]-OVTA (50pM) in the presence of 50μM unlabelled (Thr4, Gly7)-oxytocin (Bachem, Germany).

#### µ-opioid receptor binding

MOPr autoradiography was carried out in accordance with^48^. For the determination of total binding, slides were incubated for 60 minutes in 4 nM [^3^H]-tyrosyl-3,5-3H(N) (DAMGO) (PerkinElmer, 1905.5 GBq/mmol) in Tris-HCl (pH 7.4, room temperature). Adjacent sections were incubated in [^3^H]DAMGO (4nM) in the presence of 1μM naloxone (Sigma-Aldrich, UK), to determine non-specific binding (NSB).

#### mGlu_5_ receptor binding

Autoradiographic binding for mGlu_5_ receptor was carried out as previously described ^28^. For the determination of total binding, slides were incubated for 60 minutes in 10 nM [^3^H]-2-methyl-6-([3,5-3H] phenylethynl) pyridine ([^3^H]MPEP) (American Radiolabeled Chemical, 2.22 TBq/mmol) in Tris-HCl buffer (pH 7.4, 4^◦^C). Adjacent brain section were incubated with [^3^H]MPEP (10 nM) in the presence of 10μM fenobam (Tocris Bioscience, Bristol, UK), to determine the non-specific binding (NSB).

#### TSPO protein binding

TSPO autoradiography was carried out as previously described ^34^ with minor modifications. For the determination of total binding, slides were incubated for 90 minutes in 6nM [^3^H] PRB28 in Tris-HCl buffer (pH 7.4, room temperature). Adjacent brain section were incubated with [^3^H] PRB28 (6nM) in the presence of 10μM PK11195, to determine the NSB.

Slides for the OTR mGlu5R, MOPr and TSPO binding were apposed for 3 days, 3 weeks, 10 weeks 8 weeks, respectively. Slides with brains sections from all the treatment groups were laid down against the same film (Kodak BioMax MR-1 films; Sigma-Aldrich, Gillingham, UK) along with appropriate ^3^H (for MPEP, DAMGO and TSPO binding) and ^14^C (for OVTA binding) microscale standards (Amersham Pharmacia Biotech, Buckinghamshire, UK) to allow quantification, developed and analysed in parallel in a complete paired protocol. Analysis was performed using an image analyzer (MCID; Image Research, Linton, UK).

### Bioinformatics analysis via Gene Expression Omnibus (GEO)

A bioinformatics analysis was conducted to investigate the expression of metabotropic glutamate receptor 5 (mGluR5) in brain regions associated with chronic ethanol exposure. Human gene expression data set were explored using the Gene Expression Omnibus (GEO)^49^ database. Search terms included “alcohol” and brain regions of interest, including the hippocampus, nucleus accumbens, amygdala, cerebellum, and putamen. The study type was filtered to “expression profiling by array” in *Homo sapiens*.

Only human postmortem brain transcriptomic datasets comparing individuals with alcohol use disorder (or alcohol dependence/chronic alcoholism) and healthy controls were included. Analyses were performed separately for each brain region, and datasets were not pooled. The identified data sets were analyzed using GEO2R tool to compare gene expression difference between alcohol-exposed and control groups. To specifically assess mGluR5 involvement, the following search terms were used: grm5, mglur5, mglu5, glu5, and metabotropic glutamate receptor 5. A total of four datasets meeting our search criteria were identified with the following accession numbers: GSE29555, GSE62699 (platform GPL571), GSE44456, and GSE180722. The GSE180722 dataset included postmortem brain tissue from 16 male individuals, comprising 8 patients with alcohol use disorder (AUD) and 8 healthy controls. AUD diagnoses were established according to the Diagnostic and Statistical Manual of Mental Disorders, Fourth Edition (DSM-IV). The GSE29555 dataset included postmortem basolateral, central, and medial amygdala samples from 17 individuals with alcohol dependence (15 males and 2 females) and 15 healthy male controls. The GSE62699 (platform GPL571) dataset comprised postmortem nucleus accumbens tissue from 36 individuals (18 with alcohol dependence and 18 healthy controls). Cases with infectious diseases, including HIV/AIDS, hepatitis B or C, or Creutzfeldt–Jakob disease, were excluded. The GSE44456 dataset included postmortem hippocampal samples from 39 individuals (19 with chronic alcoholism and 20 healthy controls), comprising 13 males and 6 females in the alcoholic group and 13 males and 7 females in the control group.

### Statistical analysis

All the values are expressed as the mean ± SEM and have been normalized as percent change from control. All statistical analyses were performed using GraphPad v8 (GraphPad software Inc., La Jolla, CA, USA). Mixed-effects ANOVA was performed for the body weight and food intake with factors ‘treatment’ (control/ethanol) and ‘days’. Repeated-measures two-way ANOVA was performed for the locomotor activity, withdrawal symptoms, and rotarod analysis with factors ‘treatment’ (control/ethanol) and ‘time’ or ‘trial’ (repeated factor). Two-way ANOVA was performed for the NOR analysis with factors ‘treatment’ (control/ethanol) and ‘object’ (novel vs familiar). Two-way ANOVA was performed in each individual brain region with factors ‘treatment’ (control/ethanol) and ‘experimental phase’ (chronic, 1, 4 and 7 days withdrawal). ANOVAs were followed by Holm-Sidak *post-hoc* comparison if there was an interaction (statistical trend or statistical significance). The data were tested for normality fitting using QQ plots and for variance using the Brown-Forsythe test. All relevant statistical information, sample sizes are provided in **Tables 1 and 2**.

**Table 1:**
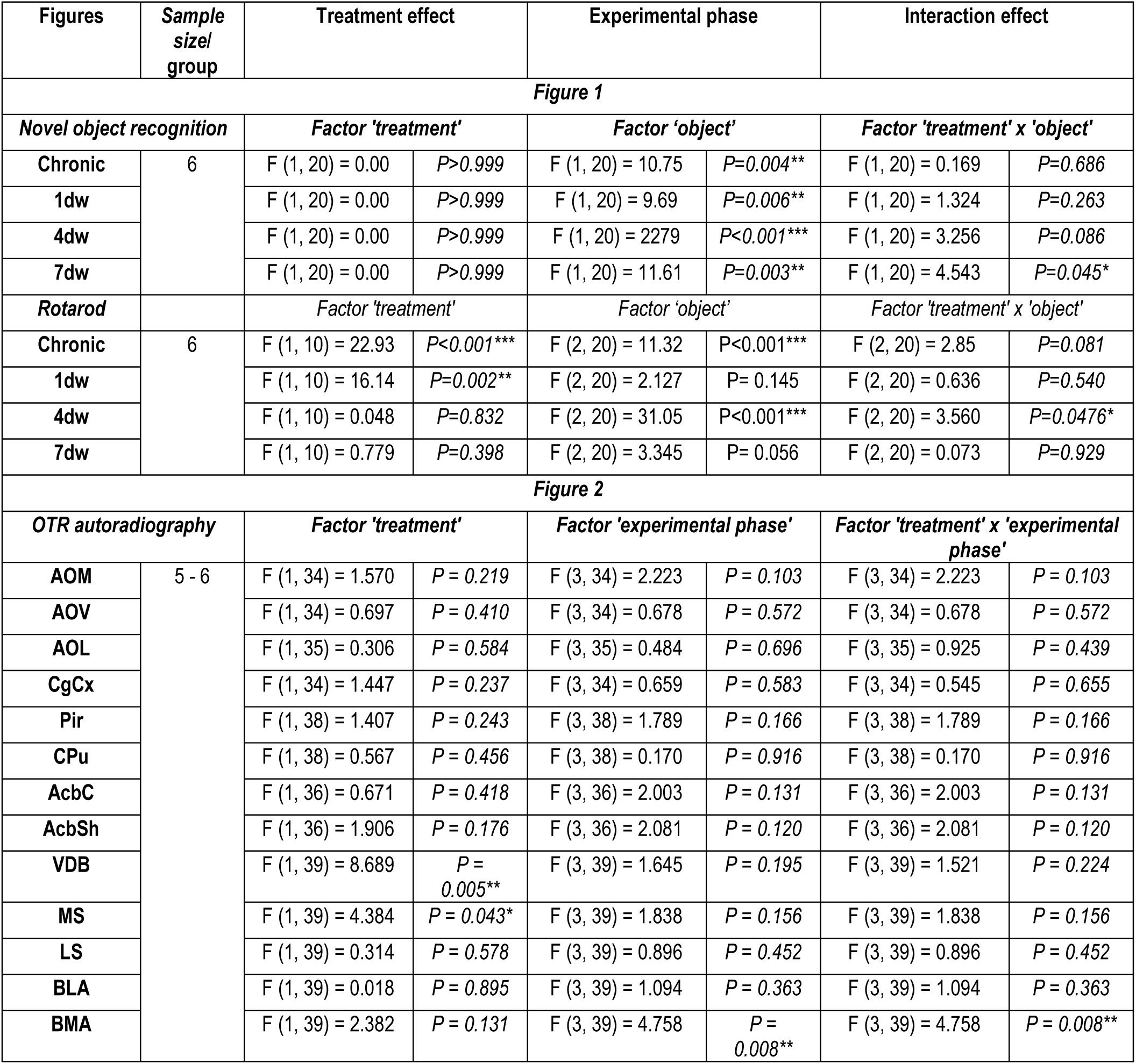
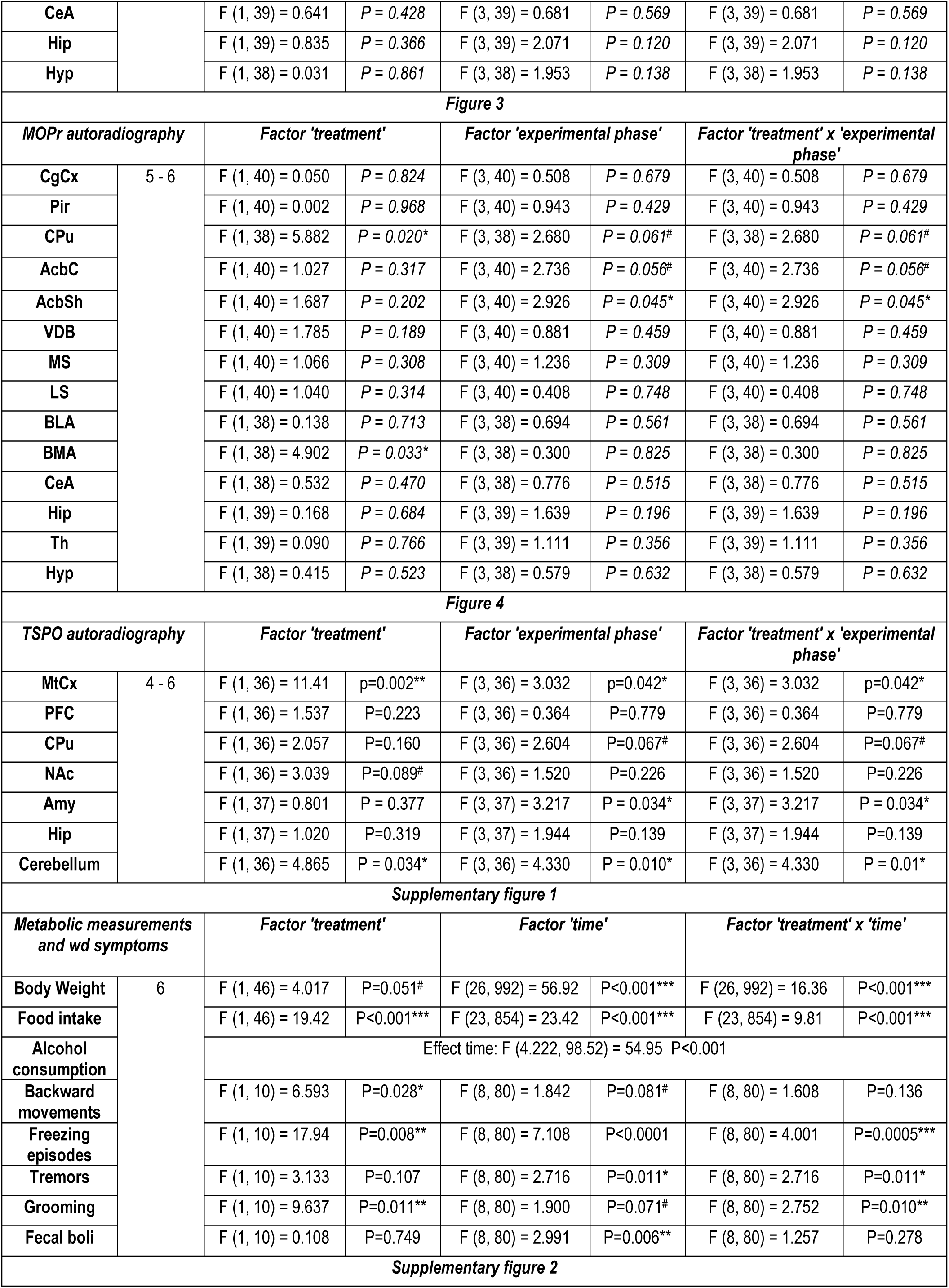
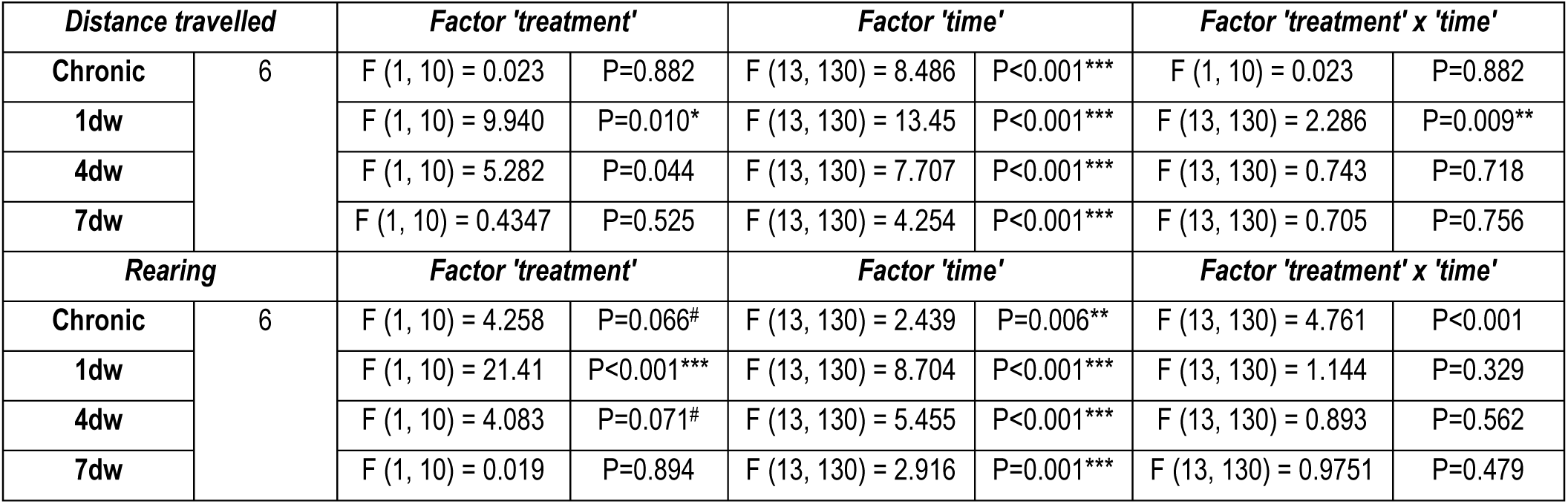
Summary of statistical analyses in the ANOVA tests.

**Table 2:** mGluR5 receptor binding following alcohol administration and withdrawal.

| [<sup>3</sup>H] MPEP binding (fmol/mg tissue) |  |  |  |  |  |  |  |  |  |  |  |  |  |  |  |  | Statistical analyses |  |  |  |
| --- | --- | --- | --- | --- | --- | --- | --- | --- | --- | --- | --- | --- | --- | --- | --- | --- | --- | --- | --- | --- |
| Brain region | Chronic |  |  |  | 1-day withdrawal |  |  |  | 4-days withdrawal |  |  |  | 7-days withdrawal |  |  |  |  |  |  |  |
|  | Control |  | Alcohol |  | Control |  | Alcohol |  | Control |  | Alcohol |  | Control |  | Alcohol |  |  |  |  |  |
|  | Mean | SEM | Mean | SEM | Mean | SEM | Mean | SEM | Mean | SEM | Mean | SEM | Mean | SEM | Mean | SEM | N | Treatment | Phase | Interaction |
| CgCx | 58.37 | 9.348 | 79.52 | 14.2 | 94.85 | 13.12 | 105.1 | 20.61 | 86.89 | 15.18 | 105.7 | 17.17 | 75.01 | 15.3 | 76.03 | 15.59 | 6,6,6,6,4,5,6,6 | F<sub>(1, 30)</sub> = 0.773; p=0.39 | F<sub>(2, 30)</sub> = 2.357; p=0.11 | F<sub>(2, 30)</sub> = 0.223; p=0.80 |
| CPu | 68.92 | 5.702 | 92.24 | 12.87 | 98.57 | 10.55 | 95.61 | 9.589 | 92.58 | 1.286 | 99.77 | 14.55 | 67.49 | 13.61 | 82.45 | 18.32 | 6,6,6,6,4,5,6,6 | F<sub>(1, 30)</sub> = 1.353; p=0.25 | F<sub>(2, 30)</sub> = 1.721; p=0.20 | F<sub>(2, 30)</sub> = 0.586; p=0.56 |
| AcbC | 71.52 | 9.026 | 104.4 | 18.18 | 94.74 | 6.097 | 122.4 | 10.8 | 95.08 | 17.94 | 104.9 | 15.2 | 80.21 | 15.36 | 93.75 | 14.64 | 6,6,6,6,4,5,6,6 | F<sub>(1, 30)</sub> = 5.399; p=0.03* | F<sub>(2, 30)</sub> = 1.753; p=0.19 | F<sub>(2, 30)</sub> = 0.296; p=0.75 |
| AcbSh | 69.27 | 7.29 | 81.05 | 17.15 | 84.27 | 11.3 | 106.5 | 12.87 | 71.37 | 13.51 | 98.17 | 16.45 | 63.76 | 15.93 | 82.5 | 14.17 | 6,6,6,6,4,5,6,6 | F<sub>(1, 30)</sub> = 2.542; p=0.12 | F<sub>(2, 30)</sub> = 1.660; p=0.21 | F<sub>(2, 30)</sub> = 0.078; p=0.93 |
| VDB | 55.65 | 9.364 | 69.11 | 20.38 | 77.04 | 13.3 | 85.99 | 18.47 | 74.49 | 13.87 | 82.17 | 19.42 | 60.09 | 16.45 | 65.54 | 16.21 | 6,6,6,6,4,5,6,6 | F<sub>(1, 30)</sub> = 0.499; p=0.49 | F<sub>(2, 30)</sub> = 0.921; p=0.41 | F<sub>(2, 30)</sub> = 0.031; p=0.97 |
| MS | 51.19 | 9.753 | 64.52 | 19.25 | 71.19 | 15.95 | 69.88 | 17.04 | 70.89 | 6.509 | 87.29 | 10.66 | 69.72 | 17.37 | 66.18 | 14.94 | 6,6,6,6,4,5,6,6 | F<sub>(1, 30)</sub> = 0.047; p=0.83 | F<sub>(2, 30)</sub> = 0.351; p=0.71 | F<sub>(2, 30)</sub> = 0.164; p=0.85 |
| LS | 86.22 | 18.65 | 129 | 23.28 | 127.9 | 14.72 | 141.4 | 18.49 | 122.5 | 15.38 | 146.5 | 16 | 100.1 | 16.35 | 99.3 | 22.92 | 6,6,6,6,4,5,6,6 | F<sub>(1, 30)</sub> = 1.377; p=0.25 | F<sub>(2, 30)</sub> = 1.800; p=0.18 | F<sub>(2, 30)</sub> = 0.661; p=0.52 |
| BLA | 77.28 | 11.11 | 91.34 | 7.385 | 68.99 | 13.18 | 103.4 | 6.231 | 96.55 | 9.69 | 92.14 | 11.98 | 59.32 | 9.977 | 85.73 | 12.33 | 6,6,6,6,6,6,6,6 | F<sub>(1, 40)</sub> = 5.649; p=0.02* | F<sub>(3, 40)</sub> = 1.475; p=0.24 | F<sub>(3, 40)</sub> = 1.301; p=0.29 |
| BMA | 76.13 | 13.8 | 75.25 | 12.07 | 66.92 | 13.85 | 86.91 | 9.554 | 84.62 | 8.823 | 78.01 | 17.41 | 66.74 | 11.53 | 81.21 | 5.415 | 6,6,6,6,6,6,6,6 | F<sub>(1, 40)</sub> = 0.626; p=0.43 | F<sub>(3, 40)</sub> = 0.135; p=0.94 | F<sub>(3, 40)</sub> = 0.541; p=0.66 |
| CeA | 60.43 | 12.74 | 86.79 | 16.23 | 68.06 | 9.706 | 85.71 | 18.3 | 73.9 | 11.28 | 69.49 | 8.663 | 60.67 | 5.883 | 88.8 | 22.42 | 6,6,6,6,6,6,6,6 | F<sub>(1, 40)</sub> = 2.878; p=0.10 | F<sub>(3, 40)</sub> = 0.047; p=0.99 | F<sub>(3, 40)</sub> = 0.560; p=0.64 |
| Hip | 84.06 | 13.23 | 109.5 | 7.173 | 85.61 | 15.45 | 97.45 | 12.21 | 86.21 | 14.45 | 91.84 | 6.021 | 71.7 | 9.923 | 96.24 | 9.618 | 6,6,6,6,6,6,6,6 | F<sub>(1, 40)</sub> = 4.332; p=0.04* | F<sub>(3, 40)</sub> = 0.433; p=0.73 | F<sub>(3, 40)</sub> = 0.361; p=0.78 |
| Th | 41.5 | 12.31 | 52.65 | 5.329 | 54.55 | 6.04 | 56.23 | 9.886 | 47.67 | 11.25 | 58.37 | 6.582 | 39.17 | 6.121 | 43.49 | 11.96 | 6,6,5,6,6,6,6,6 | F<sub>(1, 39)</sub> = 1.132; p=0.29 | F<sub>(3, 39)</sub> = 0.923; p=0.44 | F<sub>(3, 39)</sub> = 0.127; p=0.94 |
| Hyp | 60.55 | 12.26 | 74.32 | 4.801 | 58.84 | 12.82 | 66.58 | 11.65 | 50.62 | 11.98 | 65.13 | 9.333 | 45.17 | 8.642 | 57.36 | 14.55 | 6,6,6,6,6,6,6,6 | F<sub>(1, 40)</sub> = 2.347; p=0.13 | F<sub>(3, 40)</sub> = 0.771; p=0.52 | F<sub>(3, 40)</sub> = 0.037; p=0.99 |

To examine associations between behavioral performance and neurobiological measures, multiple linear regression analyses were performed with behavioral outcomes (NOR and rotarod) as the dependent variables and receptor binding measures (MOPr, OTR, mGluR5, and TSPO) as independent variables. Due to limited sample sizes within individual treatment groups, data from control and ethanol-exposed animals were combined for these exploratory analyses. Regression coefficients (β), 95% confidence intervals (CI), and corresponding p-values were calculated. Results were visualized using coefficient plots displaying β estimates and their 95% confidence intervals (**Figure 5**). This exploratory analysis as performed in only in the regions that post-hoc analysis revealed differences in any binding and were available in all the different measures.

Pearson correlation matrix analysis was performed followed by FDR for multiple comparison correction to test any significant correlations between the different bindings only in the regions that post-hoc analysis revealed differences in any binding and were available in all the different measures (**Figure 6**). The r values and significances are designated in (**Figure 6**).

For the bioinformatics analysis, differential gene expression between alcohol-dependent individuals and healthy controls was assessed separately for each dataset using the GEO2R online analysis tool (NCBI GEO), which implements the limma (Linear Models for Microarray Data) package in R. GRM5 expression was extracted from each dataset, and log2 fold change (logFC), moderated t-statistics, raw p-values, and Benjamini–Hochberg false discovery rate (FDR)-adjusted p-values were used to evaluate differential expression. Analyses were performed independently for each brain region, and no meta-analysis or pooling of datasets was conducted.

## Results

### Effect of chronic ethanol administration and one, four and seven days of withdrawal on body weight, food consumption, ethanol consumption and withdrawal symptoms

Repeated measures two-way ANOVA and pos-hoc analysis when appropriate revealed a significant effect of ethanol consumption (**Table 1**). Specifically, alcohol exposed mice failed to gain weight (**Supplementary Fig 1A**) and reduced their food intake (**Supplementary Fig 1B**) compared to control, both of which were restored short after withdrawal from alcohol. Moreover, ethanol consumption increases with the increased in the percentage provided in the liquid diet (**Supplementary Fig 1C**). Following the removal of ethanol from the mice’s diet, we observed an overall increase in backward movements (**Supplementary Fig 1D**), increase in freezing episodes following 2-,4 and 12-hours of withdrawal (**Supplementary Fig 1E**), increase in tremor episodes 4 and 8 hours post-withdrawal (**Supplementary Fig 1F**), decrease in the number of grooming 4 and 12-hours post-withdrawal (**Supplementary Fig 1G**) and no difference in the number of fecal boli (**Supplementary Fig 1H**). These symptoms disappeared following 16hrs of ethanol withdrawal and were comparable to controls.

### Effect of chronic ethanol administration and one, four and seven days of withdrawal on novel object recognition, rotarod and locomotor activity

#### Novel object recognition

Two-way ANOVA revealed an overall object effect following chronic ethanol exposure, 1 and 4-days of withdrawal (**Figure 1A-C**), suggesting intact memory recognition. An interaction effect was observed in the two-way ANOVA following 7 days of withdrawal. Holm-Sidak post-hoc analysis demonstrated that while control mice showed a significant preference to the novel object, ethanol withdrawn mice did not show any preference to either the novel or familiar object (**Figure 1D**), indicative of an impaired memory recognition.

#### Rotarod

Repeated measures two-way ANOVA revealed treatment effect following chronic exposure to ethanol (**Figure 1E**) at 1-day withdrawal (**Figure 1F**) with ethanol-treated animals performing worst. No difference between treatments was observed following 4- and 7-days of withdrawal (**Figure 1G-H**). A significant trial effect, i.e. improvement in the latency to fall between first to last trial, was observed following chronic ethanol exposure (**Figure 1E**), 4-day (**Figure 1G**) and 7-days (**Figure 1H**) but not at 1-day withdrawal (**Figure 1F**).

#### Distanced travelled

Repeated measures two-way ANOVA demonstrated an overall significant treatment effect of ethanol in the at the chronic administration (increase), 1-day (decrease) and 4 days (increase) withdrawal. Holm-Sidak post-hoc test specifically demonstrated that ethanol treatment increases the distance travelled at the 12 hours mark point, ie the first 2 hours of the dark cycle in the chronic ethanol treatment group (**Supplementary Figure 2A**). *Post-hoc* analysis also demonstrated decreased distance travelled in the 1-day withdrawal group at 8-18 hours post ethanol removal (**Supplementary Figure 2B**). An overall treatment effects was observed at 4-days withdrawal with ethanol-treated animals demonstrated higher distance travelled than control mice (**Supplementary Figure 2C**). No difference was observed at the 7-day withdrawal point (Supplementary Figure 2D).

#### Rearing

Repeated measures two-way ANOVA demonstrated an overall significant treatment effect of ethanol at the chronic administration (decrease), 1-day withdrawal (decrease). Holm-Sidak revealed a decrease in rearing during chronic exposure in ethanol-treated *vs* control animals during the dark cycle point (**Supplementary Figure 2E**). An overall lower rearing was observed in ethanol withdrawn mice at 1-day withdrawal (Supplementary Figure 2F). No difference in rearing was observed following 4- (**Supplementary Figure 2G**) and 7-days (**Supplementary Figure 2H**) of withdrawal.

### Effect of chronic ethanol administration and one, four and seven days of withdrawal on oxytocin receptor binding

Two-way ANOVA revealed a significant treatment effect in the ventral limp of diagonal band of Broca (VDB) and medial septum (MS) (**Figure 2 and Table 1**). An overall upregulation of OTR was observed in the ethanol-treated groups in these two brain regions. A significant “experimental phase” and “treatment” x “experimental phase” interaction effect was observed in the basomedial amygdala (BMA; **Table 1**). Holm-Sidak p*ost-hoc* analysis revealed a significant increase in OTR binding in the BMA only following 7 days withdrawal from ethanol consumption compared with the respective control. No significant effects were observed in any other brain region analyzed (**Figure 2**). A significant transient decrease in the BMA was observed following 4-days withdrawal *vs* chronic ethanol-treated group, followed by a rebound increase at 7days withdrawal (**Figure 2**). OTR binding returned to levels similar to the chronic administration group following the 7-days of withdrawal in the BMA (**Figure 1**).

**Figure 2:**
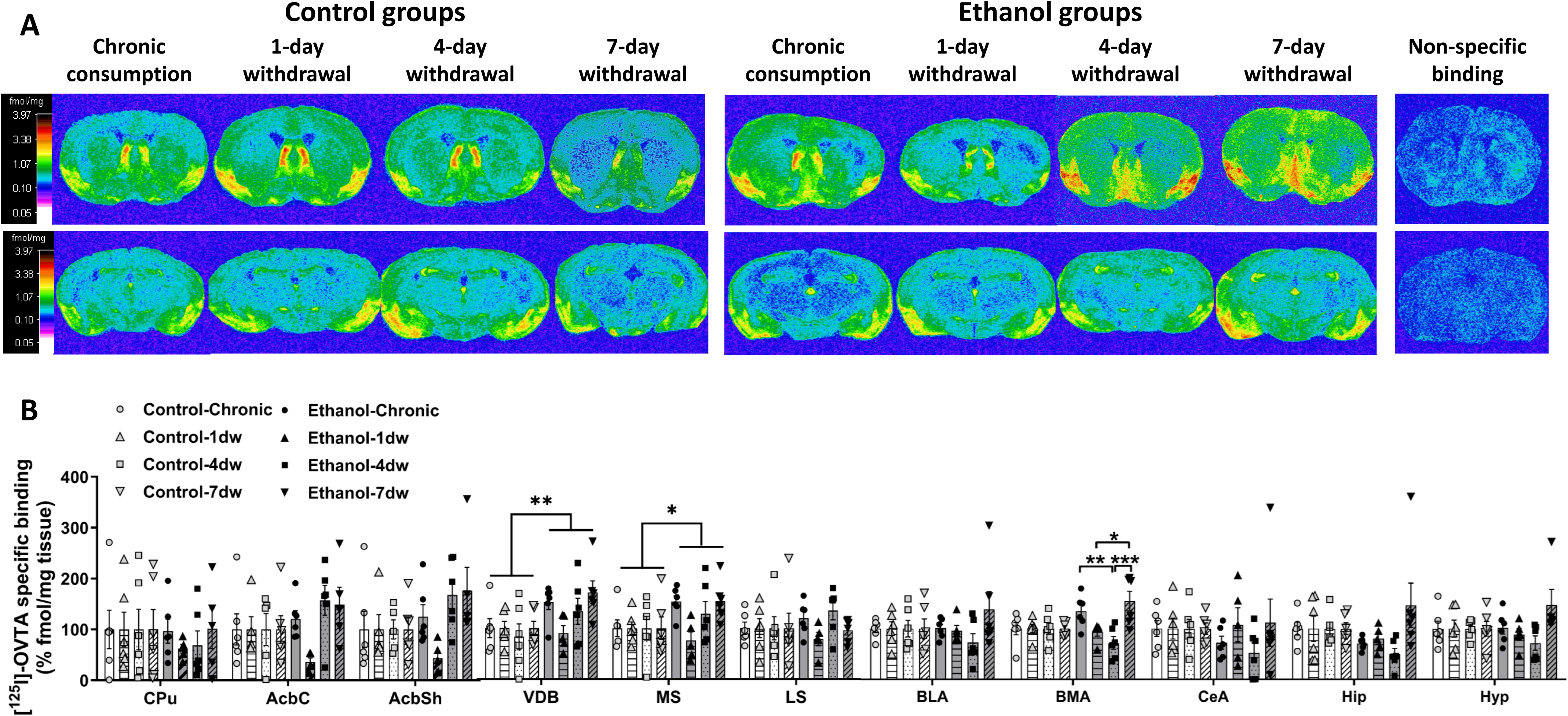
Effects of chronic ethanol consumption, acute (1-day), mid-range (4-day) and chronic (7-day) withdrawal from ethanol on the oxytocin receptor binding. Male C57BL/6J mice consumed ethanol containing diet for 10 days and then allowed to spontaneously withdraw for one, four and seven days. **(A)** Computer-enhanced representative OTR autoradiograms of adjacent coronal brain sections from chronic ethanol, acute (1-day), mid-range (4-day) and chronic (7-day) withdrawal from ethanol and their respective controls at the level of the striatum (Bregma 0.86 mm, first row) and the <u>a</u>mygdala (Bregma −2.06 mm, second row). OTRs were labelled with [^125^I]-OVTA (50pM). The color bar illustrates a pseudo-color interpretation of black and white film images in fmol/mg tissue equivalent. **(B)** Quantitative oxytocin receptor autoradiographic binding from the level of striatum, septum and forebrain from chronic ethanol consumption and withdrawal groups and their respective controls. All data are expressed as mean ± SEM (n = 5-6/group; % control). \**p <* 0.05, *p < 0.01*.

### Effect of chronic ethanol administration and one, four and seven days of withdrawal on µ-opioid receptor binding

Two-way ANOVA revealed a significant main effect of treatment in the CPu and BMA, and a significant main effect of withdrawal phase and interaction effect in the CPu, nucleus accumbens shell (AcbSh), and nucleus accumbens core (AcbC) (**Figure 3**, **Table 1**). Holm-Sidak post hoc analysis revealed a transient increase in MOPr binding following 4 days of withdrawal in the CPu, AcbC, and AcbSh vs controlswith binding levels returning to values comparable to controls after 7 days of withdrawal (**Figure 3**). In addition, a distinct temporal pattern was observed throughout the striatum, characterized by increased MOPr binding at 4 days of withdrawal compared with both the chronic alcohol treatment and 1-day withdrawal groups in the CPu, AcbC, and AcbSh. This effect was attenuated at 7 days of withdrawal relative to 4 days of withdrawal. No significant effects were observed in any other brain regions analyzed (**Table 1**).

**Figure 3:**
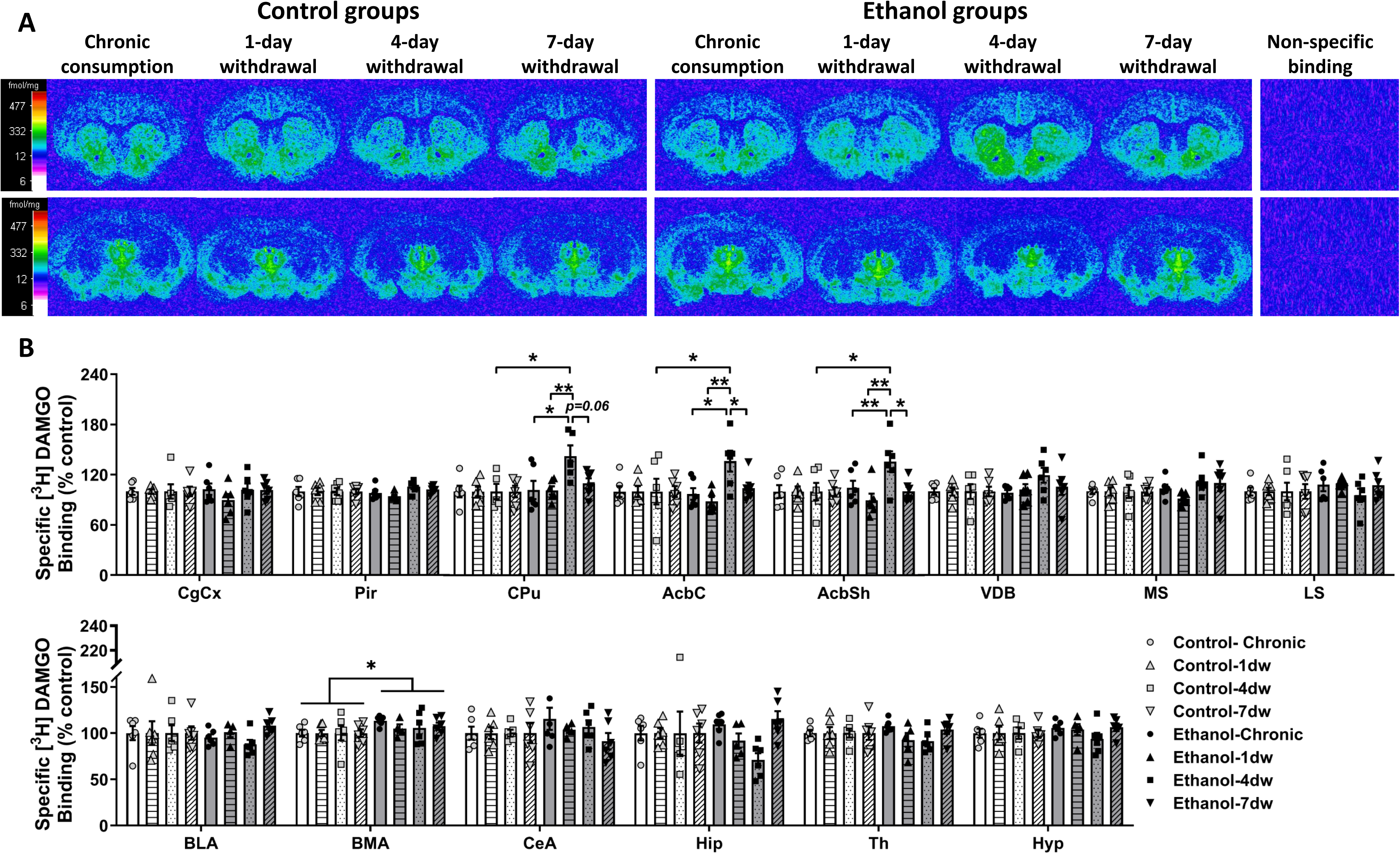
Effects of chronic ethanol consumption, acute (1-day), mid-range (4-day) and chronic (7-day) withdrawal from ethanol on the μ-opioid receptor binding. Male C57BL/6J mice consumed ethanol containing diet for 10 days and then allowed to spontaneously withdraw for one, four and seven days. **(A)** Computer-enhanced representative MOPr autoradiograms of adjacent coronal brain sections from chronic ethanol, acute (1-day), mid-range (4-day) and chronic (7-day) withdrawal from ethanol and their respective controls at the level of the <u>s</u>riatum (Bregma 0.86 mm, first row) and the amygdala (Bregma −2.06 mm, second row). MOPrs were labelled with [^3^H]-DAMGO (4nM). The color bar illustrates a pseudo-color interpretation of black and white film images in fmol/mg tissue equivalent. **(B)** Quantitative MOPr receptor autoradiographic binding from the level of olfactory, cortex, striatum, septum and forebrain from chronic ethanol consumption and withdrawal groups and their respective controls. All data are expressed as mean ± SEM (n = 5-6/group; % control). \**p <* 0.05, \*\**p* < 0.01.

### Effect of chronic ethanol administration and one, four and seven days of withdrawal on mGlu_5_ receptor binding

Two-way ANOVA revealed a significant main effect of treatment, characterized by increased mGluR5 (MPEP) binding in ethanol-treated animals compared with controls in the nucleus accumbens core (AcbC), basolateral amygdala (BLA), and hippocampus (Hip) (**Table 2**). No significant interaction was observed in any region thus *post-hoc* analysis was not performed.

### Effect of chronic ethanol administration and one, four and seven days of withdrawal on TSPO binding

Two-way ANOVA revealed a significant main effect of treatment in the motor cortex (MtCx), nucleus accumbens (NAc), and cerebellum, as well as a significant main effect of experimental phase in the MtCx, NAc, CPu, and cerebellum and a significant interaction in the MtCx, Amy, cerebellum and significant trend in the CPu (**Figure 4**, **Table 1**). In the region that a significant ANOVA interaction was found, Holm-Sidak post hoc analysis revealed a transient increase in TSPO binding in the MtCx following chronic ethanol self-administration compared with controls. TSPO binding subsequently returned to levels comparable to controls after 4, and 7 days of withdrawal (**Figure 4**). In addition, a transient increase in TSPO binding was observed after 1 day of withdrawal in the CPu, amygdala (Amy), and cerebellum compared with the corresponding control groups and the chronic ethanol group in the CPu. This increase was not maintained, with TSPO binding returning to levels comparable to controls after 7 days of withdrawal (**Figure 4**). No significant effects were observed in any other brain regions analyzed (**Table 1**).

**Figure 4:**
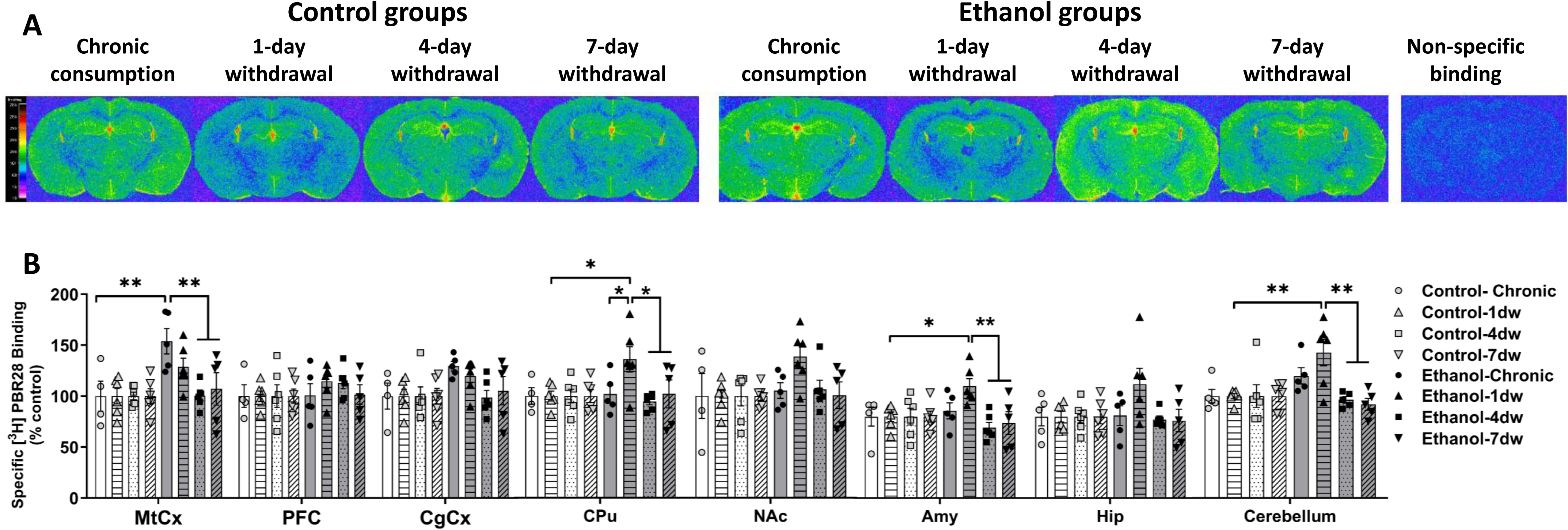
Effects of chronic ethanol consumption, acute (1-day), mid-range (4-day) and chronic (7-day) withdrawal from ethanol on the translocator protein binding. Male C57BL/6J mice consumed ethanol containing diet for 10 days and then allowed to spontaneously withdraw for one, four and seven days. **(A)** Computer-enhanced representative TSPO autoradiograms of adjacent coronal brain sections from chronic ethanol, acute (1-day), mid-range (4-day) and chronic (7-day) withdrawal from ethanol and their respective controls at the level of the forebrain (Bregma −2.06 mm). TSPO was labelled with [^3^H]-PBR28 (8nM). The color bar illustrates a pseudo-color interpretation of black and white film images in fmol/mg tissue equivalent. **(B)** Quantitative TSPO autoradiographic binding from the level of cortex, striatum, forebrain and cerebellum from chronic ethanol consumption and withdrawal groups and their respective controls. All data are expressed as mean ± SEM (n = 4-6/group; % control). \**p <* 0.05, \*\**p* < 0.01.

### Multiple linear regression analysis for relationships between behavioral and neurobiological outcomes

#### Novel object recognition

Exploratory multiple linear regression analyses revealed significant negative β coefficients for mGluR5 binding in the amygdala at 7 days of withdrawal (**Figure 5A**) and in the CPu at 1 days of withdrawal (**Figure 5A**), suggesting that higher mGluR5 binding was associated with poorer cognitive performance at this withdrawal time point. No significant relationships were identified for OTR binding (**Figure 5B**). In contrast, positive β coefficients were observed for MOPr binding in the CPu at 4 days of withdrawal (**Figure 5C**) and for TSPO binding in the CPu at 7 days of withdrawal (both p = 0.050) (**Figure 5D**), indicating that higher binding was associated with improved cognitive performance at this withdrawal time point.

**Figure 5:**
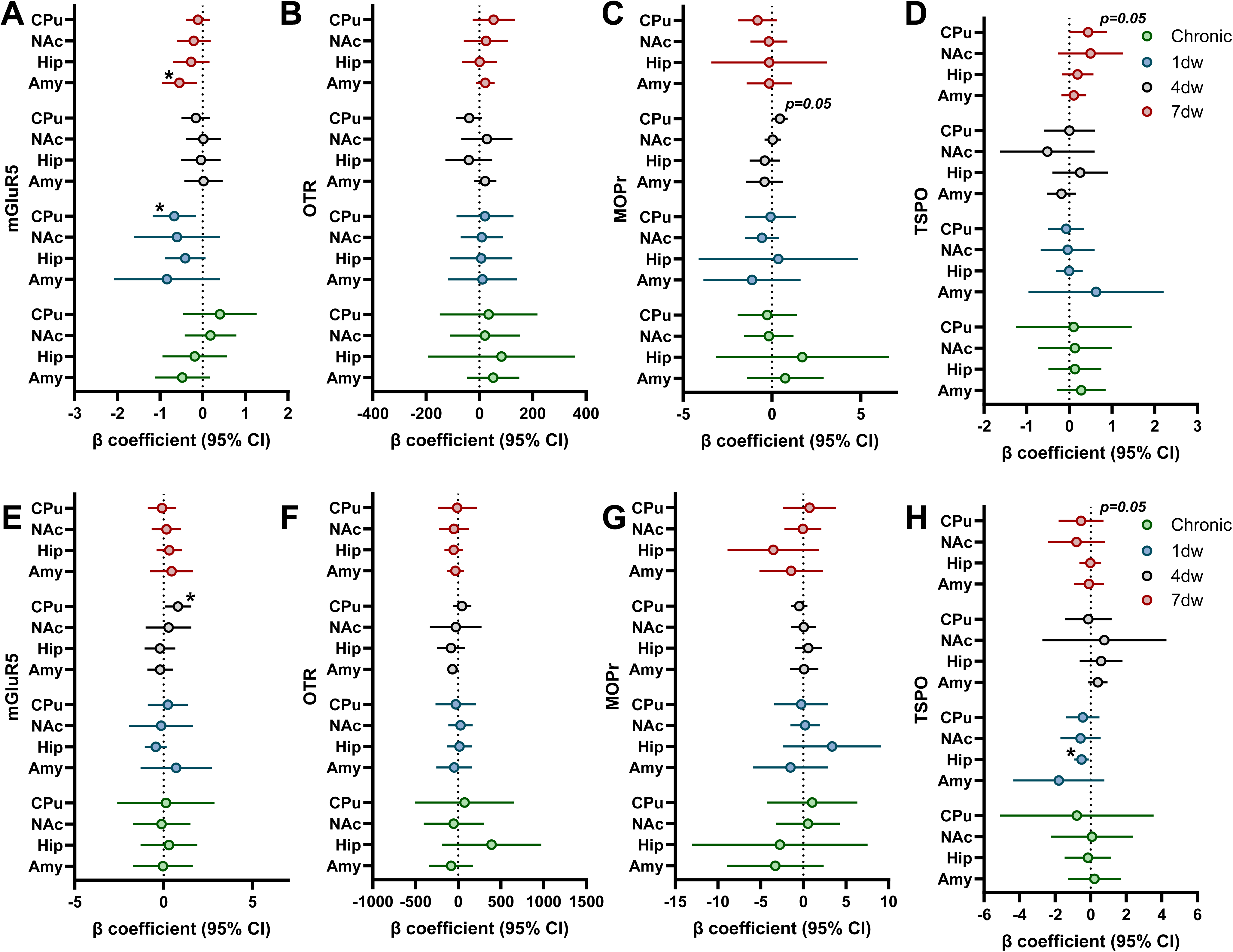
Multiple linear regression analyses examining associations between behavioral performance and receptor/transporter binding across withdrawal phases. Male C57BL/6J mice were exposed to an ethanol-containing liquid diet for 10 days and subsequently underwent 1, 4, or 7 days of withdrawal. Forest plots display β coefficients and 95% confidence intervals derived from exploratory multiple linear regression analyses assessing associations between receptor binding and behavioral performance. Novel object recognition (NOR) performance was analyzed in relation to **(A)** mGlu5R, **(B)** OTR, **(C)** MOPr, and **(D)** TSPO binding. Rotarod performance was analyzed in relation to **(E)** mGlu5R, **(F)** OTR, **(G)** MOPr, and **(H)** TSPO binding. Control and ethanol-exposed mice were combined within each withdrawal phase to maximize statistical power. Positive β coefficients indicate a positive association between receptor binding and behavioral performance, whereas negative β coefficients indicate an inverse association. Points represent β estimates and horizontal lines represent 95% confidence intervals. Significant associations are indicated by *p* < 0.05.

**Figure 6:**
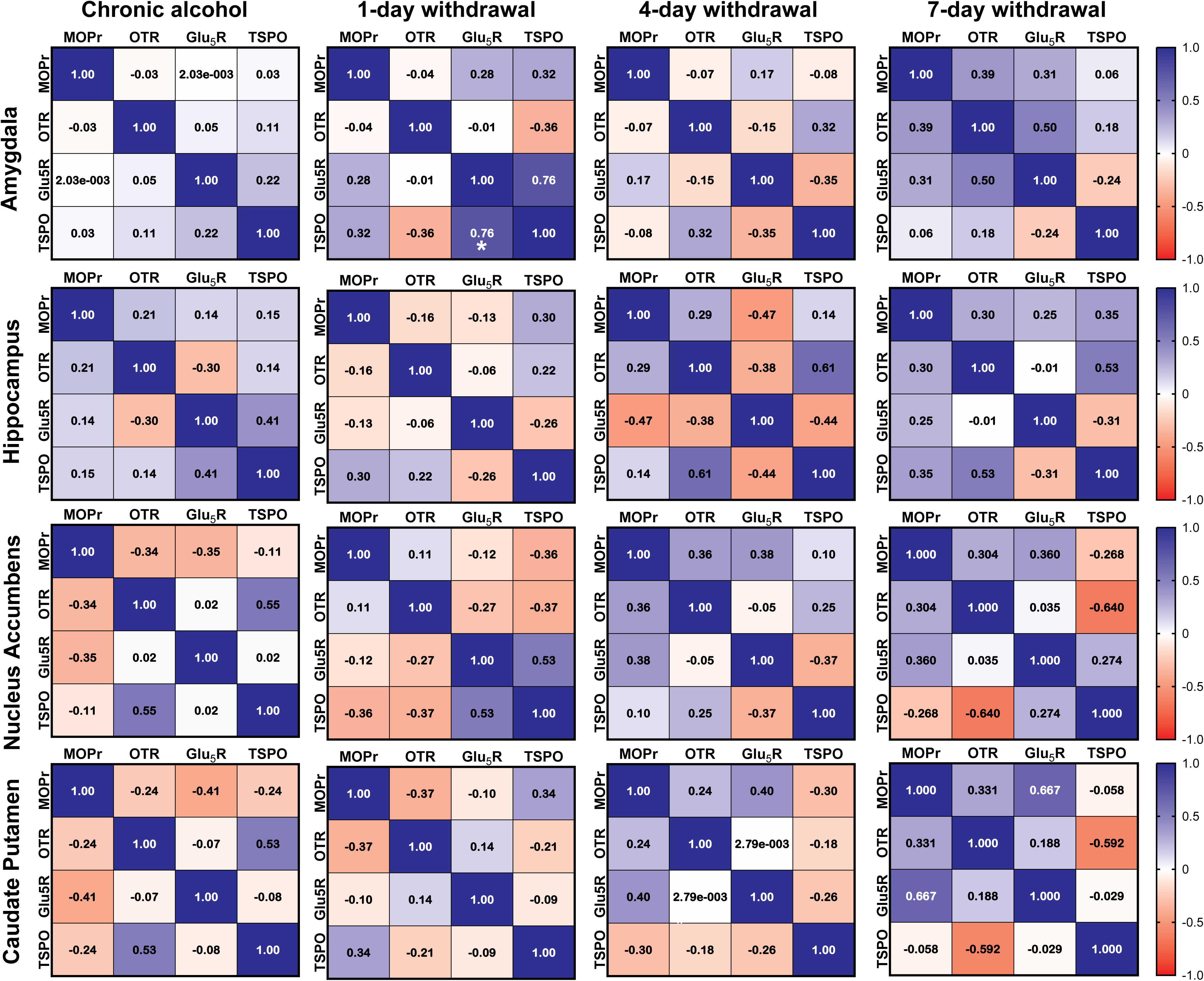
Correlation analysis between receptor/ transporter systems across ethanol withdrawal. Male C57BL/6J mice were exposed to an ethanol-containing liquid diet for 10 days and subsequently underwent 1, 4, or 7 days of withdrawal. Pearson correlation matrices were generated to examine relationships between OTR, MOPr, mGlu5R, and TSPO binding in brain regions that exhibited significant ethanol- or withdrawal-related alterations and for which all receptor measurements were available. Correlation matrices are shown fo the nucleus accumbens, caudate putamen, amygdala, and hippocampus.. Values within each matrix represent Pearson correlation coefficients for the goodness of fit (r). P-values were corrected for multiple comparisons using the Benjamini-Hochberg false discovery rate (FDR) procedure. Significant correlations following FDR correction are indicated by \**p* < 0.05.

#### Rotarod

Exploratory multiple linear regression analyses revealed a significant positive β coefficient was observed for mGluR5 binding in the CPu at 4 days of withdrawal, indicating a positive association with rotarod performance (**Figure 5E**). No significant associations were identified for OTR or MOPr binding (**Figure 5F, G**). In contrast, significant negative β coefficients were observed for TSPO binding in the CPu at 7 days of withdrawal (*p* = 0.05) and in the hippocampus at 4 days of withdrawal, suggesting that higher TSPO binding was associated with poorer rotarod performance (**Figure 5H**).

### Correlation analysis between receptor bindings and behavioral outcomes

A strong positive correlation was observed between mGluR5 and TSPO binding following 1 day of withdrawal (r = 0.76), which remained significant following FDR correction (adjusted *p* < 0.05). This suggests that increased mGluR5 binding was associated with increased TSPO binding during early withdrawal. Although several other correlations were observed between TSPO and OTR in the amygdala at 7dw and hippocampus at 4dw, as well as between mGluR5 and MPr in the CPu at 7dw they did not survive FDR correction.

### Bioinformatics analyses

Differential gene expression analysis was performed using GEO2R across all identified datasets. The analysis focused on GRM5 because mGlu5R exhibited the strongest and most consistent associations with recognition memory deficits in our preclinical experiments, making it the most translationally relevant target for validation in human postmortem datasets. However, mGluR5 (GRM5) expression was only detected in three out of the four datasets (**Table 3**).

**Table 3:** Summary of GRM5 (mGluR5) Expression Across Datasets.

| Dataset | Brain Region(s) | GRM5 Detected | Evidence of Differential Expression |
| --- | --- | --- | --- |
| GSE180722 | Amygdala | Yes | Not significant (trend only) |
|  | Cerebellum | Yes | Not significant |
|  | Hippocampus | Yes | Not significant |
|  | Putamen | Yes | Not significant |
| GSE44456 | Hippocampus | Yes | Not significant |
| GSE62699 (GPL571) | Nucleus accumbens | Yes | Inconsistent (probe-dependent; borderline significance) |
| GSE29555 | Amygdala (basolateral, central, medial nuclei) | No | Not detected among DEGs |

In the GSE180722 dataset (amygdala; **Table 3**), GRM5 (mGluR5) expression showed a moderate increase in individuals diagnosed with alcohol use disorders and actively still drinking compared to controls (logFC = 0.46). However, this difference did not reach statistical significance (*p* = 0.065, adj. *p* = 1.00). The relatively low t-statistic (*t* = 1.95) and negative B-value (−4.18) indicate weak evidence for differential expression. In the GSE180722 dataset (cerebellum; **Table 3**), GRM5 (mGluR5) expression showed no significant difference between individuals exposed to chronic alcohol use and controls (logFC = 0.09, *p* = 0.622, adj. *p* = 1.00). The low t-statistic (*t* = 0.50) and negative B-value (−4.83) indicate very weak evidence for differential expression. In the GSE180722 dataset (hippocampus; **Table 3**), GRM5 (mGluR5) expression showed no significant difference between individuals exposed to chronic alcohol use and controls (logFC = 0.26, *p* = 0.246, adj. *p* = 0.875). The low t-statistic (*t* = 1.20) and negative B-value (−4.56) indicate weak evidence for differential expression. In the GSE180722 dataset (putamen; **Table 3**), GRM5 (mGluR5) expression showed no significant difference between individuals exposed to chronic alcohol use and controls (logFC = 0.19, *p* = 0.437, adj. *p* = 1.00). The low t-statistic (*t* = 0.80) and negative B-value (−4.73) indicate weak evidence for differential expression. Overall, the GSE180722 dataset (**Table 3**) shows no statistically significant or consistent changes in GRM5 (mGluR5) expression across brain regions in individuals exposed to chronic alcohol use.

In the GSE44456 dataset (**Table 3**), differential expression analysis showed no significant difference in GRM5 expression in the hippocampus between individuals exposed to chronic alcohol use and controls (logFC = 0.13, *p* = 0.421, adj. *p* = 0.8465). The low t-statistic (*t* = 0.81) and negative B-value (−5.34) further indicate weak evidence for differential expression. These results do not provide evidence that GRM5 mRNA expression is altered in this dataset.

In the GSE62699 dataset (**Table 3**) (platform GPL571), one GRM5 probe (214217_at) showed higher expression in the nucleus accumbens of individuals exposed to chronic alcohol use compared to controls (logFC = 0.74), with a nominally significant *p*-value (*p* = 0.0043) but a non-significant adjusted *p*-value (adj. *p* = 0.0516). However, a second GRM5 probe (207235_s_at) showed minimal change (logFC = 0.05) and was not statistically significant (p = 0.265, adj. *p* = 0.523). This inconsistency between probes suggests that evidence for differential GRM5 expression in this dataset is limited.

In the GSE29555 dataset (**Table 3**), GRM5 (mGluR5) was not identified among the differentially expressed genes in the basolateral, central, and medial nuclei of the amygdala.

Overall, GRM5 was detected in most datasets but showed no consistent or statistically significant differential expression across brain regions, with only limited and inconsistent evidence observed in the nucleus accumbens.

## Discussion

In the present study, we demonstrate brain region-specific and time-dependent alterations in OTR, MOPr, mGlu5R, and TSPO binding following chronic ethanol exposure and withdrawal. These changes occur across distinct phases of withdrawal and are accompanied by the emergence of behavioral impairments. Importantly, the temporal pattern of behavioral changes observed in this model mirrors key clinical features of alcohol withdrawal and abstinence. Physical withdrawal symptoms and motor impairments were most prominent during acute withdrawal and resolved with prolonged abstinence, whereas recognition memory deficits emerged only during protracted withdrawal. Similar dissociations have been reported in individuals with alcohol use disorder, in whom acute somatic withdrawal symptoms often subside within days while cognitive deficits may persist for weeks or months following detoxification. Furthermore, the absence of locomotor and motor coordination deficits at 7 days of withdrawal indicates that the impaired performance in the novel object recognition task is unlikely to reflect reduced exploration, sedation, or generalized motor dysfunction, and instead supports a deficit in recognition memory. Together, these findings highlight the importance of considering withdrawal as a dynamic process characterized by temporally distinct neuroadaptations, which may require different therapeutic strategies depending on the stage of abstinence.

Within this model, we observed increased OTR binding in the medial septum and ventral limb of the diagonal band of Broca following ethanol exposure and withdrawal. These regions are critically involved in the regulation of stress and affective processing, and septal oxytocin signaling has been implicated in behavioral responses to drugs of abuse ^25,45,47,50,51^. The observed upregulation may therefore reflect a compensatory mechanism engaged to counteract withdrawal-related stress and negative affect. A similar pattern of septal OTR upregulation has been reported following exposure to other addictive substances, supporting the notion of a shared neuroadaptive response across drug classes ^25,45,47,50,51^. In addition, we identified a decrease of OTR binding in the basomedial amygdala at 4 days withdrawal that was restored after 7 days of withdrawal. The temporal pattern of OTR regulation in the basomedial amygdala suggests that oxytocinergic adaptations may be more relevant to the evolution of withdrawal states rather than acute withdrawal symptoms per se. Specifically, OTR binding showed a transient reduction during early withdrawal, a period characterized by evident physical withdrawal symptoms and heightened physiological stress, followed by a rebound increase after 7 days of withdrawal. One possible interpretation is that the delayed upregulation represents a compensatory neuroadaptation aimed at counteracting the negative affective and stress-related consequences of prolonged abstinence. This interpretation is consistent with the established role of oxytocin signaling in stress regulation and the ability of oxytocin to attenuate withdrawal-related behaviors in both preclinical and clinical studies. Although we did not observe a direct association between amygdalar OTR binding and recognition memory or motor performance, the present findings suggest that oxytocinergic adaptations are unlikely to be directly associated with the behavioral outcomes examined in this study. Whether these alterations contribute to withdrawal-related stress- or affective-related processes remains to be determined.

Alterations in the opioidergic system followed a distinct temporal pattern during ethanol withdrawal. In agreement with previous preclinical and clinical studies demonstrating dysregulation of the endogenous opioid system following chronic alcohol exposure^52–57^, we observed an overall effect of ethanol treatment on MOPr binding in the CPu and basomedial amygdala. More notably, striatal MOPr binding exhibited a transient increase during intermediate withdrawal, peaking at 4 days of withdrawal in the CPu, AcbC and AcbSh before returning toward baseline levels by 7 days. This temporal profile may reflect a compensatory response to reduced endogenous opioid tone, as chronic ethanol exposure has been associated with decreased β-endorphin signaling^58–65^. Given the established role of striatal MOPr signaling in reward processing and motivational regulation, this transient upregulation may represent a neuroadaptive response occurring during intermediate withdrawal. Furthermore, exploratory regression analyses revealed a positive association between MOPr binding and recognition memory performance in the CPu at 4 days of withdrawal, indicating that higher MOPr binding was associated with better recognition memory performance during this withdrawal phase. However, these findings are correlational and do not establish a causal role for MOPr signaling in cognitive function. Whether elevated MOPr binding reflects a compensatory mechanism that protects against withdrawal-associated cognitive impairment remains to be determined. The transient nature of these alterations suggests that opioidergic adaptations may play a stage-specific role during abstinence rather than reflecting a persistent consequence of ethanol exposure. Such temporal regulation may have therapeutic implications and suggests that the efficacy of interventions targeting the endogenous opioid system could depend on the stage of withdrawal.

Among all neurobiological measures examined, mGlu5R exhibited the most consistent relationship with behavioral outcomes. Although ethanol exposure produced a general increase in mGlu5R binding in the nucleus accumbens core, basolateral amygdala, and hippocampus, the most notable finding was the association between mGlu5R binding and recognition memory performance. Exploratory regression analyses revealed significant negative β coefficients between mGlu5R binding and novel object recognition performance in both the amygdala (7dw) and CPu (1dw), suggesting that elevated mGlu5R availability was associated with poorer cognitive outcomes. Given the well-established role of mGlu5 receptors in synaptic plasticity^66^ and learning-related processes^67^, these findings suggest that the association between mGlu5R signaling and recognition memory is both withdrawal phase- and region-specific. Specifically, mGlu5R binding in the CPu was associated with recognition memory performance during early withdrawal, whereas amygdalar mGlu5R binding was associated with recognition memory during protracted withdrawal. These findings suggest that the neurobiological substrates underlying memory function may shift across withdrawal stages. Early withdrawal may depend more heavily on striatal adaptations, whereas during protracted withdrawal amygdalar glutamatergic signaling may become increasingly relevant to cognitive function. Furthermore, mGlu5R was the only receptor system to show consistent associations with both recognition memory and motor learning measures, emphasizing its potential importance in mediating withdrawal-related behavioral dysfunction. The observed association between mGlu5R binding and rotarod performance further suggests that glutamatergic adaptations may influence both cognitive and motor domains during withdrawal, although these relationships appear to depend on the stage of abstinence. The translational relevance of this finding is strengthened by growing evidence implicating mGlu5R in alcohol use disorder^68^ and supports further investigation of mGlu5R as a potential target for alleviating alcohol withdrawal-associated cognitive impairments.

Alongside these neurotransmitter-specific adaptations, ethanol exposure and withdrawal produced transient alterations in TSPO binding, indicative of dynamic neuroimmune responses across the withdrawal period. Chronic ethanol exposure increased TSPO binding in the motor cortex, whereas acute withdrawal was characterized by transient elevations in TSPO binding within the caudate-putamen, amygdala, and cerebellum that returned to control-like levels following prolonged abstinence. This temporal profile is consistent with the view that neuroimmune activation represents an early response to ethanol withdrawal rather than a persistent feature of protracted abstinence^69^. Although associations between TSPO binding and behavioral measures were relatively limited, exploratory regression analyses identified region- and time-specific relationships with both recognition memory and motor performance. Specifically, higher TSPO binding in the CPu during protracted withdrawal was associated with improved recognition memory performance, whereas increased TSPO binding in the CPu and hippocampus was associated with poorer rotarod performance during protracted and acute withdrawal stages, respectively. Although associations between TSPO binding and behavioral measures were identified, their interpretation remains unclear and may depend on both withdrawal stage and brain region. Interestingly, TSPO binding positively correlated with mGlu5R binding during early withdrawal, suggesting that neuroimmune and glutamatergic adaptations may occur in parallel during this phase of abstinence. In line with our findings, acute immune stressors such as lipopolysaccharide-induced neuroinflammation have been shown to induce simultaneous upregulation of both TSPO and mGlu5R^70^. While the causal relationship between these adaptations cannot be determined from the present study, these findings raise the possibility that early neuroimmune activation may contribute to, or occur alongside, glutamatergic adaptations during ethanol withdrawal. The observation that TSPO alterations were most evident during acute withdrawal further suggests that neuroimmune processes may represent an early and potentially time-sensitive target for therapeutic intervention.

Taken together, these findings demonstrate that ethanol withdrawal is characterized by temporally distinct neuroadaptations across multiple neurobiological systems. Acute withdrawal was associated with transient increases in TSPO binding, suggesting an early neuroimmune response, whereas intermediate withdrawal was characterized by transient alterations in striatal MOPr and amygdala OTR binding. In contrast, mGlu5R alterations showed the strongest and most consistent relationship with behavioral outcomes at specific withdrawal time points, particularly recognition memory performance. The delayed emergence of recognition memory deficits during protracted withdrawal, together with the observed associations between mGlu5R binding and cognitive performance, suggests that glutamatergic dysregulation may contribute to withdrawal-associated cognitive dysfunction. In addition, the positive correlation between mGlu5R and TSPO binding during early withdrawal supports the possibility that glutamatergic and neuroimmune adaptations evolve in parallel during abstinence. The absence of consistent GRM5 transcriptional alterations in human postmortem datasets further suggests that receptor-level changes may not necessarily be reflected by changes in gene expression and may instead involve post-transcriptional mechanisms, altered receptor trafficking, post-translational regulation, or other factors not captured by transcriptomic analyses.

Several limitations should be considered when interpreting these findings. First, although significant associations were identified between receptor binding and behavioral outcomes, the regression analyses were exploratory in nature and performed in a relatively small sample, warranting replication in larger cohorts. Second, the present study was designed to characterize temporal neurobiological adaptations during ethanol withdrawal and therefore does not establish causal relationships between receptor alterations and behavioral impairments. Future studies using pharmacological or genetic manipulations will be required to determine whether the observed changes in mGlu5R, MOPr, OTR, or TSPO signaling directly contribute to withdrawal-associated behavioral deficits. Finally, the use of male mice exclusively limits the generalizability of these findings, and future work should examine whether similar temporal adaptations occur in females.

In conclusion, chronic ethanol exposure and withdrawal produce region-specific and temporally distinct alterations in oxytocinergic, opioidergic, glutamatergic, and neuroimmune systems. These adaptations are accompanied by behavioral impairments that emerge at different stages of withdrawal, with mGlu5R showing the strongest and most consistent association with recognition memory deficits. Together, these findings highlight withdrawal as a dynamic process characterized by stage-specific neuroadaptations and suggest that distinct therapeutic strategies may be required across abstinence, including targeting neuroimmune processes during acute withdrawal and glutamatergic dysfunction during protracted abstinence.

## Acknowledgments

This study was supported by Royal Society grant (RG120556, PI: Alexis Bailey) and Brain and Behavior Research foundation grant (NARSAD young investigator award, PI: Polymnia Georgiou). The sponsors had no involvement in the design of the study and in the collection, analyses and interpretation of the data, nor in the writing of the report and the decision to submit this article for publication.

## Author contributions

*Designed research:* P.Georgiou, L. Wells and A. Bailey; *Performed research:* P. Georgiou, S.A. Zimmerman, A. Onisiforou, F. Pantouli, P. Zanos, M. Sklirou; *Analyzed data:* P. Georgiou, J.A. Garcia-Carmona, F. Pantouli*; Wrote or contributed to the writing of the manuscript:* All authors

## Data Accessibility

Primary data and material information are provided in the manuscript. If further information is required, please contact the corresponding author.

---

**Supplementary Figure 1: Effects of chronic ethanol consumption, acute (1-day), mid-range (4-day) and chronic (7-day) withdrawal from ethanol on body weight, food intake and withdrawal symptoms.** Male C57BL/6J mice consumed ethanol containing diet for 10 days and then allowed to spontaneously withdraw for one, four and seven days. **(A)** Body weight and **(B)** food intake prior the introduction of ethanol in the mice’s liquid diet, during the 10-day ethanol consumption and during the 7-day withdrawal. **(C)** Ethanol consumption during the 10-day administration period. **(D-H)** Withdrawal symptoms scored following 2-96 hours if withdrawal. All data are expressed as mean ± SEM (n = 6/group). \**p <* 0.05, \*\**p < 0.01*, *\*\*\*p < 0.001*.

**Supplementary Figure 2: Effects of chronic ethanol consumption, acute (1-day), mid-range (4-day) and chronic (7-day) withdrawal from ethanol on locomotor activity.** Male C57BL/6J mice consumed ethanol containing diet for 10 days and then allowed to spontaneously withdraw for one, four and seven days. Distance travelled **(A)** during the last day of chronic ethanol exposure and following **(B)** 1-day, **(C)** 4-days and **(D)** 7-days of withdrawal. Rearing frequency (E) during the last day of chronic ethanol exposure and following **(F)** 1-day, **(G)** 4-days and **(H)** 7-days of withdrawal. All data are expressed as mean ± SEM (n = 6/group). \**p <* 0.05, \*\**p<0.01*, *\*\*\*p < 0.001*.

