## Supplementary figures and images for "Ethanol withdrawal induces temporally distinct alterations in oxytocinergic, opioidergic, glutamatergic and neuroimmune systems associated with recognition memory deficits"

### Supplementary Fig 1

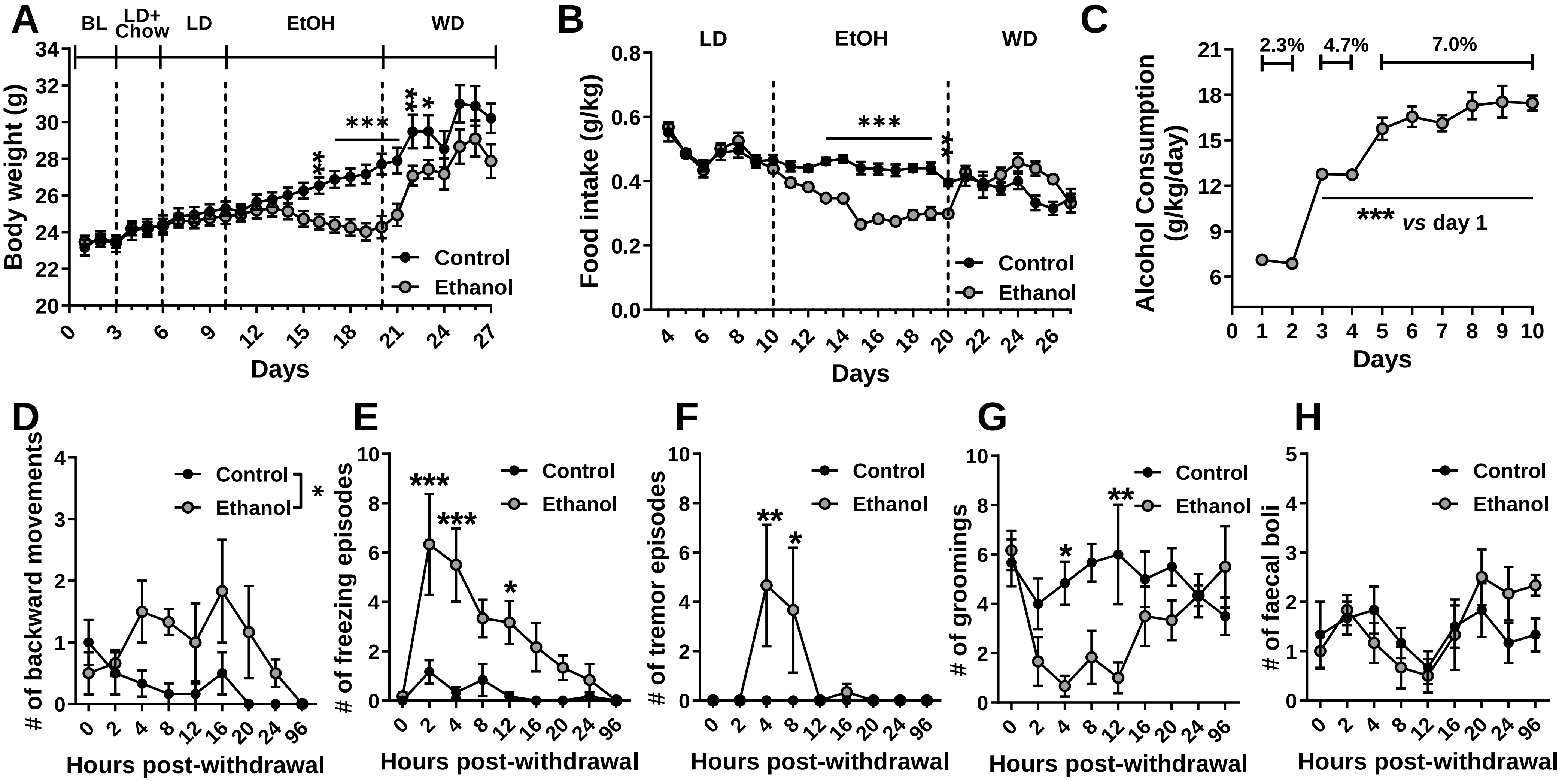

### Supplementary Fig 2

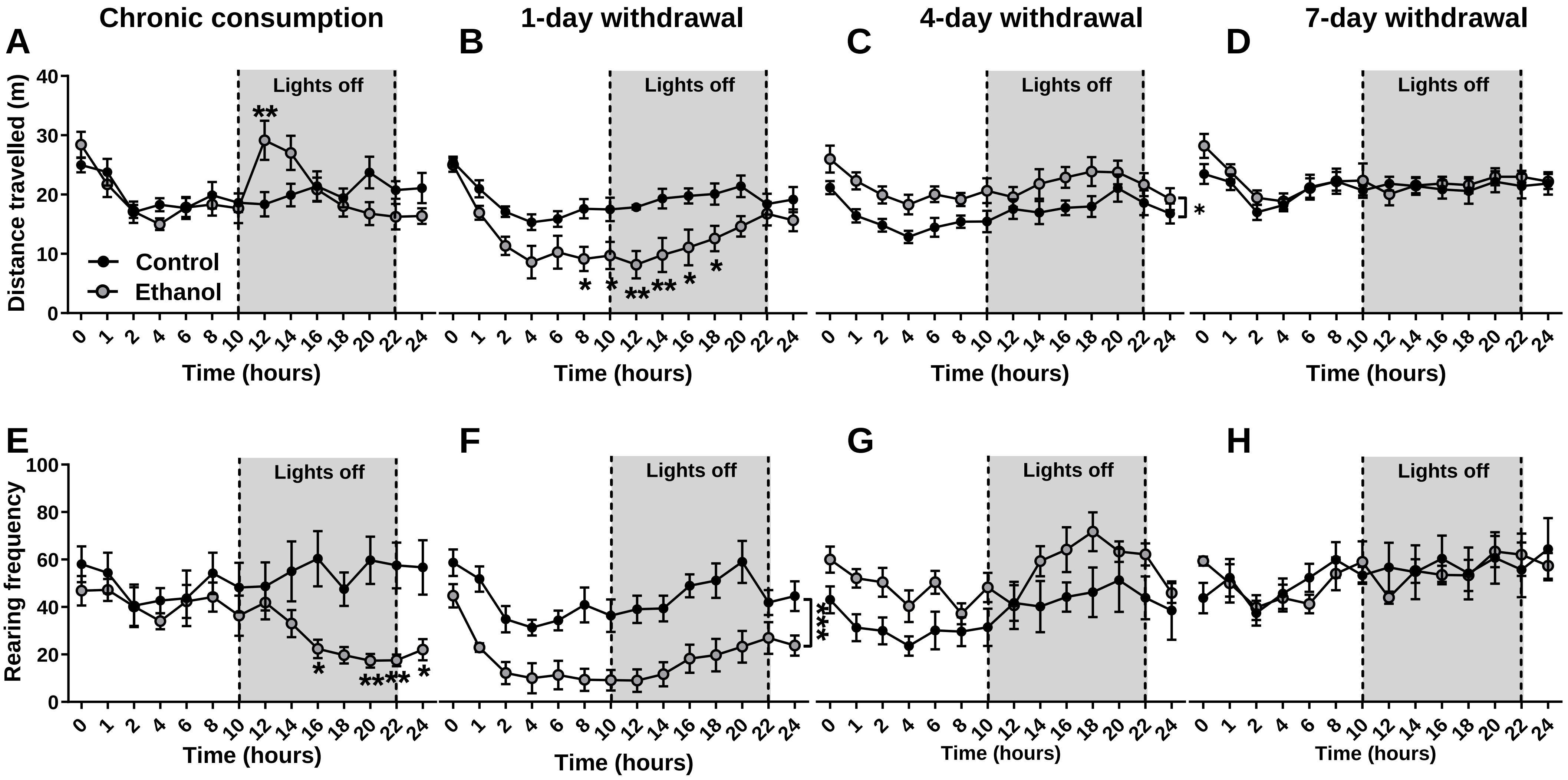
